# The balance hypothesis for the avian lumbosacral organ and an exploration of its morphological variation

**DOI:** 10.1101/2020.04.01.020982

**Authors:** Kathryn E. Stanchak, Cooper French, David J. Perkel, Bingni W. Brunton

## Abstract

Birds (Aves) exhibit exceptional and diverse locomotor behaviors, including the exquisite ability to balance on two feet. How birds so precisely control their movements may be partly explained by a set of intriguing modifications in their lower spine. These modifications are collectively known as the lumbosacral organ (LSO) and are found in the fused lumbosacral vertebrae called the synsacrum. They include a set of transverse canal-like recesses in the synsacrum that align with lateral lobes of the spinal cord, as well as a dorsal groove in the spinal cord that houses an egg-shaped glycogen body. Based on compelling but primarily observational data, the most recent functional hypotheses for the LSO consider it to be a secondary balance organ, in which the transverse canals are analogous to the semicircular canals of the inner ear. If correct, this hypothesis would reshape our understanding of avian locomotion, yet the LSO has been largely overlooked in the recent literature. Here, we review the current evidence for this hypothesis and then explore a possible relationship between the LSO and balance-intensive locomotor ecologies. Our comparative morphological dataset consists of micro-computed tomography (μ-CT) scans of synsacra from ecologically diverse species. We find that birds that perch tend to have more prominent transverse canals, suggesting that the LSO is useful for balance-intensive behaviors. We then identify the crucial outstanding questions about LSO structure and function. The LSO may be a key innovation that allows independent but coordinated motion of the head and the body, and a full understanding of its function and evolution will require multiple interdisciplinary research efforts.

**Jargon-Free:** Birds have an uncanny ability to move their heads independently of their bodies. They can keep their heads remarkably still to focus on their prey while twisting in flight or perched on a bouncing branch. How are they able to do this? Like us, birds have balance organs in their inner ears that act like gyroscopes. Surprisingly, birds may have an additional balance organ known as the “lumbosacral organ” in their spine, right above their legs, which might help them sense the movement of their body separately from their head. This second balance organ may have played a very important role in bird evolution and how birds move, but it has seldom been considered in recent scientific studies. This intriguing hypothesis is based in part on a series of fluid-filled, canal-like recesses in the bone surrounding the spinal cord, which resemble the semicircular canals of the inner ear. We looked for evidence of these canal-like recesses in many different bird species, and we found it in every bird we examined. We also found that birds that perch often have deeper recesses than birds that do not perch, suggesting these canals help maintain balance. This paper presents those findings, reviews the existing research, and identifies some key questions that need to be asked to advance our understanding of this fascinating and mysterious part of the bird spinal cord.

## INTRODUCTION

Over evolutionary time, land-dwelling vertebrates have co-opted their limbs to perform a variety of tasks. Birds (Aves) have accumulated a particularly impressive set of locomotor specializations, including bipedalism, flight, and the superb ability to balance on a perch. To facilitate the use of limbs, the spinal columns of tetrapods have evolved many morphological and neural specializations. These include expansions of the spinal cord that support sensory processing and motor control of the fore- and hindlimbs and that originate from the brachial and lumbar/sacral plexuses, respectively. Birds have a particularly prominent posterior expansion housed within the synsacrum (Fig. 1A, Giffin, 1990), a bony fusion of lumbar and sacral vertebrae that are themselves fused to the pelvic bones (Baumel and Witmer, 1993). In the avian synsacrum, the vertebral canal--which houses the spinal cord--greatly expands in diameter, more so than in other diapsids (Giffin, 1990). This synsacral vertebral canal expansion houses a set of unique anatomical features collectively known as the avian “lumbosacral organ” or LSO (Figs. 1B and 2).

**Figure 1.**
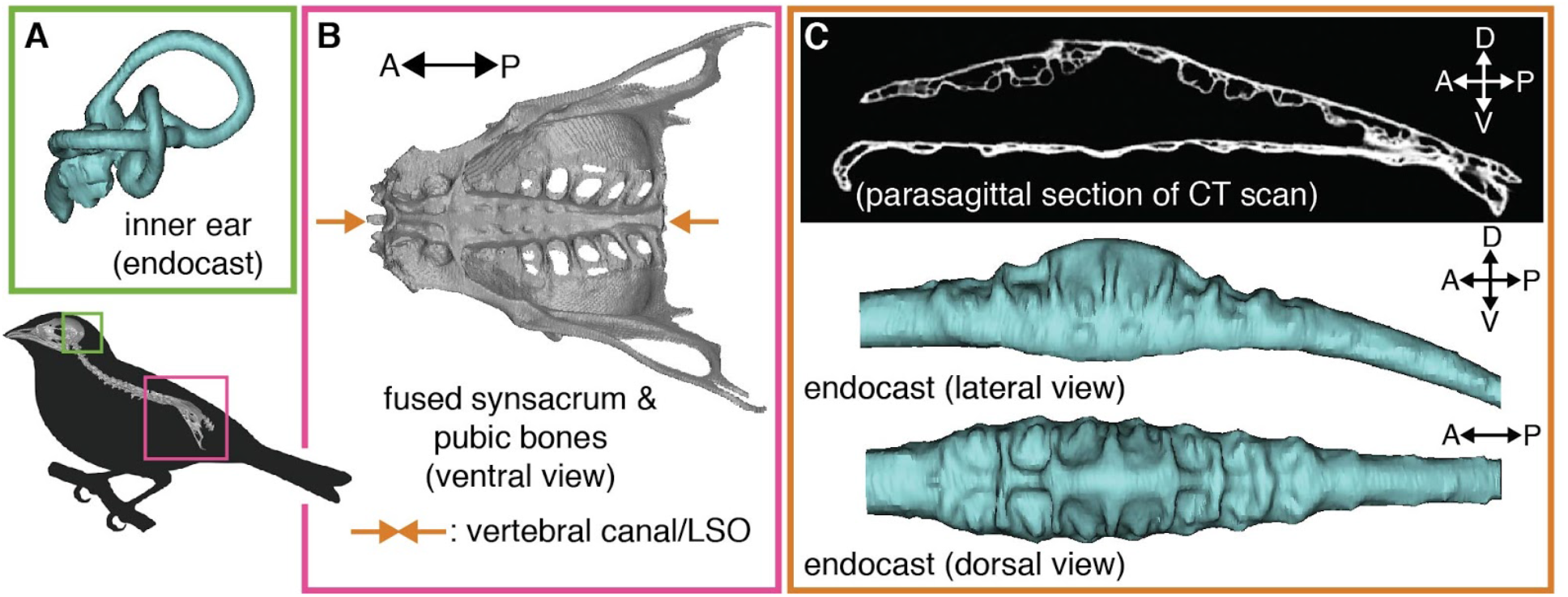
Birds are hypothesized to have two sets of balance organs, one in the inner ear (A) (common to terrestrial vertebrates) and a second within the synsacrum (B), termed the lumbosacral organ (LSO). The LSO is housed within an expanded vertebral canal, which exhibits a set of canal-like recesses (lumbosacral transverse canals, or LSTCs; C).

**Figure 2.**
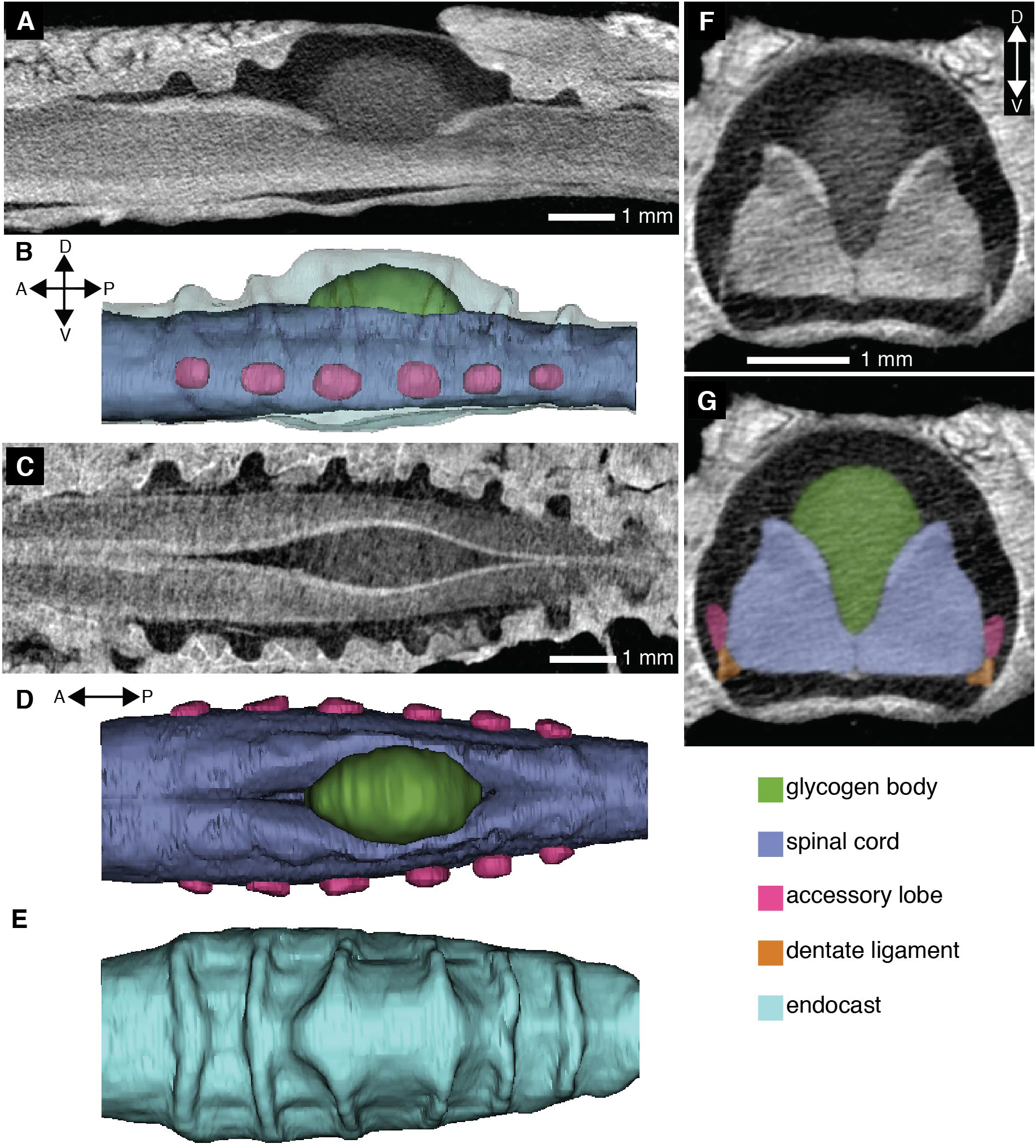
A contrast-enhanced CT scan demonstrates the soft tissue morphology of the avian lumbosacral organ (LSO) and the space within the lumbosacral transverse canals (LSTCs). A parasagittal section (A) shows the LSTCs and the expansion in the vertebral canal for the glycogen body. A corresponding lateral view of a 3D model from the CT scan demonstrates the accessory lobes (B). A horizontal section (C) and the corresponding dorsal view of the soft tissue in the model (D) show the glycogen body within the dorsal groove of the spinal cord. Ridges evident in a dorsal view of the endocast of the vertebral canal (E) result from the LSTCs. A transverse section of the third canal from the anterior end (F) is paired with labeled soft tissue elements (G).

The LSO is distinct from other parts of the avian spinal column in both its osteology and its soft tissue morphology. Specifically, the LSO consists of both a specialized vertebral canal morphology in the synsacrum and a distinctive lumbosacral spinal cord (Figs. 1 and 2). The synsacral vertebral canal exhibits a series of transverse recesses along its dorsum (Necker, 1999); these resemble canals in shape, so here we refer to them as lumbosacral transverse canals (LSTCs). In addition, the spinal cord splits dorsally, along the sagittal plane, and within this split rests an ovoid organ of glycogen-rich cells called the glycogen body (De Gennaro, 1982). The spinal cord in this region is supported by thickened dentate ligaments as well as transverse ligaments (Schroeder and Murray, 1987). In addition to the LSTCs, the splitting of the spinal cord, and the glycogen body, the LSO also includes a set of accessory lobes that project laterally from the ventrolateral aspect of the lumbosacral spinal cord. These accessory lobes are pairs of spinal cord protrusions that align with the transverse canals (Necker, 1999).

Several hypotheses have been put forward for the function of the LSO and its various components, but detailed understanding of this remarkable specialization remains elusive. For instance, a frequent—and perhaps the most obvious—functional hypothesis for the glycogen body is that it is used for energy storage, but this hypothesis is not supported by all experimental evidence (summarized in De Gennaro, 1982). More recently, the LSTCs and accessory lobes have been interpreted as having a mechanosensory function (summarized in Necker, 2006). Necker reasoned that rotations of the body induce flow of cerebrospinal fluid within the transverse canals, stimulating neurons of the accessory lobes (summarized in Necker, 2006). This may help birds maintain balance, although strong electrophysiological and behavioral evidence in support of this hypothesis is still needed.

In this paper, we review existing hypotheses of avian LSO function and present observations to explore the specific hypothesis that the LSO has a mechanosensory role in balance control (Necker, 2006). A key prediction of this balance organ hypothesis is that anatomical variation in the LSO corresponds with locomotor function in birds. The ability to maintain balance becomes more difficult with increasing functional requirements, like keeping the head still to track prey or predators during locomotion. Anatomical adaptations in balance organs can mitigate functional difficulties related to locomotion (e.g., Muller, 1994; Spoor et al., 2007; Grohé et al., 2018).

Using μ-CT scans of synsacra representing 44 species, 37 families and 25 orders across Aves, we investigate a possible relationship between locomotor lifestyle and LSO morphology. We found osteological evidence of the LSO (i.e., LSTCs and a vertebral canal expansion) in all bird species examined. We also found a trend of more prominent LSTCs in species that perch. Interpreted in combination with the existing literature, our data refines the hypothesis of the LSO’s function as a putative balance organ. We identify critical knowledge gaps that currently prevent a thorough understanding of the avian LSO and suggest key opportunities to further elucidate LSO function with modern tools and approaches.

## HYPOTHESES OF LSO FUNCTION

Over the past century, several lines of research have posited functional hypotheses for the various anatomical components of the LSO. The most prominent of these suggest a metabolic role for the glycogen body and a balance/mechanosensory role for the accessory lobes and LSTCs. Although these two hypotheses are not necessarily mutually exclusive, they have generally been discussed in the literature in isolation of one another.

### The glycogen body as a metabolic repository

The cells of the glycogen body are metabolically active, but there is no consistent evidence that the glycogen body has a role in body metabolism. The evidence from starvation-type studies investigating a potential role of the glycogen body in metabolism are equivocal. Many (but not all) found a reduction in glycogen in the liver without an accompanying loss within the glycogen body (as summarized in De Gennaro, 1982). Notably, enzymatic assays revealed no presence of glucose-6-phosphatase (Benzo et al., 1975; Fink et al. 1975), which is an enzyme necessary for the conversion of glucose-6-phosphate to free glucose.

While the liver uses glycogen for energy storage, with subsequent transport of glucose to other parts of the body for glycolytic conversion, the absence of glucose-6-phosphatase in the glycogen body suggests it does not share this function. Alternative metabolic roles have been posed for the glycogen body. These include that it secretes glycogen directly into cerebrospinal fluid (Azoitia et al., 1985); assists in lipid production, perhaps for myelination (Benzo et al., 1975); or that it provides lactate or pyruvate, rather than glycogen and glucose, as energy sources (Dezza et al., 1970; Fink et al. 1975).

### A mechanosensory function for the LSO

Several related biomechanical and neurophysiological mechanisms have been suggested for a potential mechanosensory role of the LSO. Schroeder and Murray (1987) suggested multiple ways in which locomotor movements might be transmitted to sensory neurons in the accessory lobes of the LSO: through tension in the dentate ligaments or meninges or through motion of the cerebrospinal fluid. Necker (2006) elaborated upon a cerebrospinal fluid-related mechanosensory role for the LSO by analogizing the LSO with the vestibular region of the inner ear. In the LSO, the LSTCs are formed by the dorsal canals in the synsacrum and closed off by the arachnoid membranes, forming proper canals that are analogous to the semicircular canals of the inner ear. According to the hypothesis, as the body experiences angular acceleration in the roll direction (i.e. around the longitudinal axis of the spinal cord) cerebrospinal fluid motion in these canals stimulates neurons in the accessory lobes.

Evidence supporting the mechanosensory hypothesis promulgated by Necker comes from neuroanatomical, physiological, and behavioral studies. If the neurons in the accessory lobes function as suggested, they must be capable of (1) sensing cerebrospinal fluid motion in the LSTCs and (2) transmitting that sensory information to motor control systems. Rosenberg and Necker (2002) found dendritic projections of the accessory lobe neurons that extend into fluid-filled lacunae, analogous to the stereocilia of inner ear hair cells. Cells in the accessory lobes have neuronal-type voltage-gated sodium channels (Yamanaka et al., 2008, Matsushita et al., 2018), indicating the presence of neurons that can generate action potentials and are thus capable of acting as Necker suggests. In addition, there is evidence that implicates GABA (Yamanaka et al., 2012), glutamate (Yamanaka et al., 2013), and acetylcholine (Takahashi et al., 2015) in accessory lobe neuron function. However, no specific transduction mechanism has been identified (as noted by Matsushita et al., 2018). Necker (1997) traced the accessory lobe neuron axons to the contralateral lamina VIII of the spinal cord, which is implicated in mammals in left-right limb coordination (Harrison et al., 1986). This connection suggests that the accessory lobes may play a role in lower limb reflexes.

Necker et al. (2000; also summarized in Necker, 2006) conducted LSO specific behavioral experiments in pigeons. They removed the dorsal bone of the synsacrum in the LSO region, thereby removing the cerebrospinal fluid and presumably preventing the stimulation of accessory lobe neurons. They reported that experimental birds could fly but had difficulty perching or walking. These behaviors were particularly disturbed in masked birds, as these birds could not use vision to compensate for their lack of balance-sensing. Interestingly, these birds appeared to try to regain balance using flight-specific reflexes instead of bipedal locomotion-specific reflexes (i.e., extending wing on the same side of the body rather than the opposite; Necker et al. 2000, Necker, 2006). Although these qualitative behavioral observations are consistent with a connection between cerebrospinal fluid and locomotion, the experiments used only a small number of individuals and do not reveal a mechanism by which the cerebrospinal fluid might interact with neurons in the spinal cord. Additional behavioral and physiological tests of the LSO balance organ hypothesis are required that include rigorously-defined experimental trials with quantitative assays, full controls, and replication.

There is also some indirect evidence for a role of the LSO in balance or bipedal locomotion. In a developmental study, Necker (2005) found that recently hatched, precocial chicks had more developed LSOs than more altricial pigeons, which might help chicks balance soon after hatching. Urbina-Meléndez et al. (2018) used a robotics approach and found that accelerometers located at the “hip” of a standing bird-like robot in addition to sensors at the “head” improved estimation of foot acceleration, which supports the idea that the LSO might help with balance.

### Mechanosensation in vertebrate spinal cords

Although more evidence is still required to definitively determine mechanosensation in the avian LSO, vertebrates are known to have mechanosensory neurons in their spinal cords that detect local mechanical stimuli. Lamprey spinal cords have a class of neurons along their lateral sides called “edge cells”—stretch receptors that respond to local strain deformation of the spinal cord (Grillner et al., 1984). These edge cells are not, however, known in other vertebrates (Henderson et al., 2019), although cells with morphologies suggestive of mechanoreception have been reported in the lateral margin of the spinal cords in some other species (e.g., reptiles, Schroeder, 1986; and amphibians, Schroeder and Egar, 1990).

Another set of sensory neurons referred to as “cerebrospinal fluid-contacting neurons,” or CSF-cNs, are found across vertebrates (Kolmer, 1921; Agduhr 1922; Orts-Del’Immagine and Wyart, 2017). CSF-cNs contact the cerebrospinal fluid in the central canal of the spinal cord. In zebrafish, they are incorporated into a spinal central pattern generator (Wyart et al., 2009) and have been shown to respond to bending of the spinal cord (Böhm et al., 2016). Further research on mechanosensory inputs to the vertebrate spinal cord are summarized in a recent review by Henderson et al. (2019).

### A comparative approach to understanding the LSO

Existing research studies are consistent with a role for the LSO in mechanosensation, particularly in response to body rotation in order to maintain balance. However, conclusive experimental studies are still lacking. Surprisingly, the LSO has received relatively little recent attention, despite the potentially critical role of the LSO in understanding avian locomotion and widespread interest in the mechanical and neural basis of sensorimotor control. One useful strategy for advancing understanding of the LSO is a comparative approach. The morphological and ecological diversity in Aves presents an opportunity for testing whether elaborated LSO morphology corresponds with balance-intensive locomotor strategies.

How the LSO varies among avian species is largely unknown. Previous descriptions of LSTC variation among birds made only qualitative comparisons (Imhof, 1905; Jelgersma, 1951; Necker, 2006). Necker (2006) concluded that these canals were relatively similar in most species examined, although he noted shallower LSTCs in a bird that almost exclusively flies (*Apus apus*, common swift) and deeper LSTCs in a bird that exclusively walks (*Struthio camelus*, ostrich). Methods for making high-resolution, quantitative observations of the internal structure of the vertebral canal have recently become commonplace (i.e., computed tomography X-ray scanning, or CT scans). Here we expand upon previous work by compiling a dataset of measurements of the synsacral vertebral canal that houses and forms part of the LSO. Using these data, we quantitatively explore the morphological disparity of the LSO across Aves, particularly in the context of diversity in locomotor ecology.

## COMPARATIVE EXPLORATION OF LSO OSTEOLOGY

### Methods

We sampled synsacra from 44 species of 25 orders from extant Aves. The majority of synsacra in our sample came from skeletal preparations of museum specimens, but two (*Taeniopygia guttata* and *Zonotrichia leucophrys*) were fluid-preserved specimens from laboratory colonies. For most species our sample consisted of one specimen, but for five species (*Apus pacificus*, *Colius striatus*, *Colaptes auratus*, *Hemiprocne mystacea*, and *Progne subis*) our sample consisted of five or six specimens. These multiple samples allowed us to assess the extent of intraspecific variation relative to interspecific variation in synsacral vertebral canal osteology. A list of all museum specimens used in this study is provided in the Supplemental Information.

We CT scanned the synsacra on either a Skyscan 1172 or 1173 (Bruker) or an NSI X5000 (North Star Imaging). For all species, we calculated two dimensionless metrics that we used to compare the morphology of the synsacral vertebral canal across species. Both metrics are based on a series of cross-sectional areas of the vertebral canal that correspond with each LSTC and inter-LSTC (intercanal) space (Fig. 2). Because the LSTCs are neither perfectly transverse nor orthogonal to the longitudinal axis of the vertebral canal, we used the CT scan viewing software Dataviewer (Bruker) to generate cross-section images that align with the LSTCs (Fig. 3). The LSTCs differ in the angle they make relative to the longitudinal axis of the neural canal, so the cross-sections differed as well. For consistency, the sets of cross-sectional images for all species were all generated by the same researcher. Then, we used a fill function in ImageJ (v. 2.0.0-rc-69/1.52p) to calculate the cross-sectional area for each image. This procedure sometimes required manually closing gaps in the cross-section image created by the spinal nerve-root foramina.

**Figure 3.**
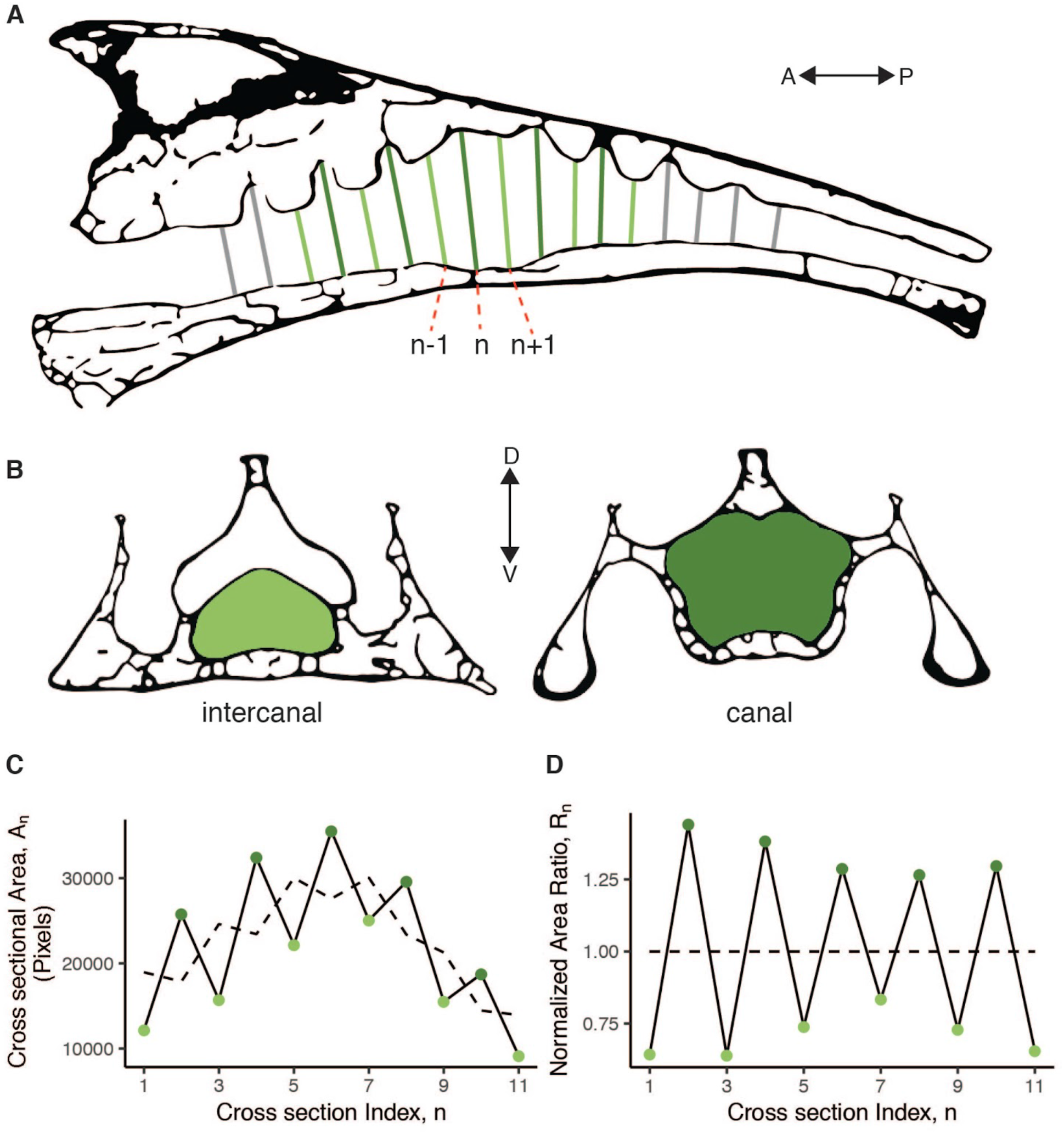
The measure of LSTC prominence is calculated from a series of cross-sectional areas from the vertebral canal. The diagram of the parasagittal section (A) illustrates how we selected cross-sections approximately perpendicular to the long axis of the vertebral canal. The labeled sections are examples corresponding with the formula for the dimensionless ratio calculation provided in the text. The transverse section diagrams (B) illustrate the cross-sectional areas of the vertebral canal for canal and intercanal sections. A plot of the cross-sectional areas is provided in (C), with a dashed line representing the running mean. A plot of the corresponding ratios is in (D). The LSTC prominence metric is defined as the standard deviation of these ratios.

The first metric we calculated, the “LSTC prominence”, is a measure of the size of the LSTCs relative to the primary vertebral canal. To calculate the LSTC prominence, we first calculated a dimensionless ratio of each cross-sectional area of that area to itself and the areas immediately anterior and posterior (see Fig. 2C and D):

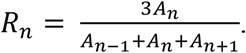

If LSTC recesses are absent, as they are in parts of the avian vertebral canal outside the synsacrum, then this ratio *R_n_* would be close to unity (1) for all n. The running mean was calculated using the “movmean” function in Matlab R2019b, which calculates a mean over a defined window of neighboring points (in this case, three). The first and last points (in this case, *n*=1 and *n*=11) have a window that falls outside of the range, so the mean at those positions is calculated using only *A_n_* and either *A_n+1_* or *A_n-1_*, respectively. To summarize across several adjacent segments, we then calculated the standard deviation of the set of ratios calculated for each synsacrum, so that the LSTC prominence is defined as the standard deviation of *R_n_* values. For many species, it was difficult to distinguish the first and last canal, as the canal depth decreases toward the anterior and posterior ends of the LSO region such that the final recess may not reasonably be considered a canal. Consequently, we constrained the number of canals considered in the LSTC prominence calculation to five, for a total of 11 cross-sectional areas: the maximum area (near the middle of the LSO region) and five sections each in the anterior and posterior directions (Fig. 3).

The second metric we calculated is a measure of the expansion of the LSO region that houses the glycogen body relative to the cross-sectional area of the spinal cord. We termed this the “expansion ratio”. Specifically, the expansion ratio is the maximum of the running mean calculated for the LSTC prominence metric divided by the mean of the most anterior cross-sectional area and the most posterior. Because we use the running mean, the expansion ratio is not influenced by transverse canal prominence.

Although selecting the cross-sectional areas is somewhat arbitrary, the LSTC prominence calculation is inherently robust to random, unbiased error in cross-sectional area measurements. In addition, we are using both calculated metrics, rather than absolute values, to consider morphological trends, so our interpretations of our results should not be affected by unbiased measurement error. Finally, for a subset of species across the morphospace, we generated endocasts of the neural canal using Mimics v.22 (Materialise) so that we could visualize 3D morphological variation in the endocasts.

We analyzed our computed metrics visually, using the packages *ape* (Paradis and Schliep, 2018), *geiger* (Harmon et al., 2008), *ggtree* (Yu et al., 2017; Yu et al., 2018), *treeio* (Wang et al., 2019), *ggplot2* (Wickham, 2016), *egg* (Auguié, 2019), *cowplot* (Wilke, 2019) and *dplyr* (Wickham et al., 2019) in R version 3.6.2 (R Core Team, 2019) to compare the LSTC prominence and expansion ratio metrics across birds with different locomotor lifestyles. We also used a genus-level phylogeny of Aves from Cooney et al., 2017 based on Jetz et al., 2012 and Prum et al., 2015 to visualize phylogenetic variation in locomotor lifestyle and LSTC prominence. To define locomotor lifestyle/mode for each species, we assigned each species a binary “yes/no” value to “bipedal” and “perch” locomotor categories. We used the recent editions of the Handbook of the Birds of the World (del Hoyo, et al., 2019) and the Birds of North America (Rodewald, ed., 2015) as well as our own collective knowledge to determine these categories. We based our assignments not on whether or not the species is capable of any particular locomotor mode, but whether or not the locomotor mode is critical to their overall lifestyle. Some species were more difficult to classify than others. For example, we classified *Colius striatus*, the spectacled mousebird, as “perching,” (which we define as balancing on a branch or other elevated, narrow object) although it might more accurately be described as “clinging.” (hanging from the side of an object). “Bipedal” refers specifically to terrestrial bipedality. Although these classifications are rudimentary and somewhat subjective, our goal was not to use them for hypothesis testing but rather for visual detection of general trends in LSO osteological morphology. To confirm visually-identified trends, we performed two-sample, two-sided Kolmogorov-Smirnov tests of our metrics using the “ks.test” function in the R *stats* package. These tests do not account for phylogenetic nonindependence of the species in our sample (see Felsenstein, 1985), and we emphasize again that we use them not to test an explicit hypothesis, but rather to explore a broadly sampled, quantitative dataset of LSO morphology, which is lacking from the existing literature.

We also conducted an exploration of the LSO soft tissue anatomy in two species of passerines present in laboratory colonies at the University of Washington: *Zonotrichia leucophrys*, the white-crowned sparrow, and *Taeniopygia guttata*, the zebra finch. The samples were incidental tissue byproducts of other experiments. Our primary interest was confirming the presence of unfilled space in the LSTCs for cerebrospinal fluid to flow. Some evidence for this has been shown histologically (Necker, 2006), but we used an alternative method, diffusible-iodine contrast-enhanced CT scanning (diceCT; Gignac et al., 2016), to assess the extent of the tissue-filled space within the lumbosacral vertebral canal. We dissected the synsacrum from one specimen of each of the two species and placed it in a 3% w/v I_2_KI solution for 4 days, scanned it in a Skyscan 1172 μ-CT scanner (Bruker), replaced it in the solution, and then rescanned it after 11 total days.

### Results and Discussion

We found osteological evidence of the LSO in the synsacra from all species we examined. Further, the morphology of the vertebral canal varied considerably among species, and this variation was evident both visually in CT scan sections and endocasts (Fig. 4) and in our calculated metrics (Fig. 5). Notable variation exists across species in the proportion of the synsacral length taken up by the LSTC region and in the number of LSTCs (e.g., *Spheniscus humboldti* vs. *Podargus strigoides* in Fig. 4). We occasionally found it difficult to determine the number of LSTCs and the extent of the LSTC region, as some species did not exhibit an obvious first or last canal. The orientation of the LSTCs relative to the longitudinal axis of the synsacral vertebral canal varies both across species and within an individual (see dorsal views in Fig. 4). If these canals serve as part of a rotational accelerometer, as Necker (2006) suggested, then this variation in canal orientation might allow sensing of motion in different directions. The shape of the canals also varies (see parasagittal views in Fig. 4). Some canals are expanded dorsally and then constrict near the vertebral canal, forming a balloon-or mushroom-like shape. The functional significance of this shape variation is unclear. The constrictions may allow the arachnoid membranes to form a more defined canal that optimizes the flow of cerebrospinal fluid toward the accessory lobes.

**Figure 4.**
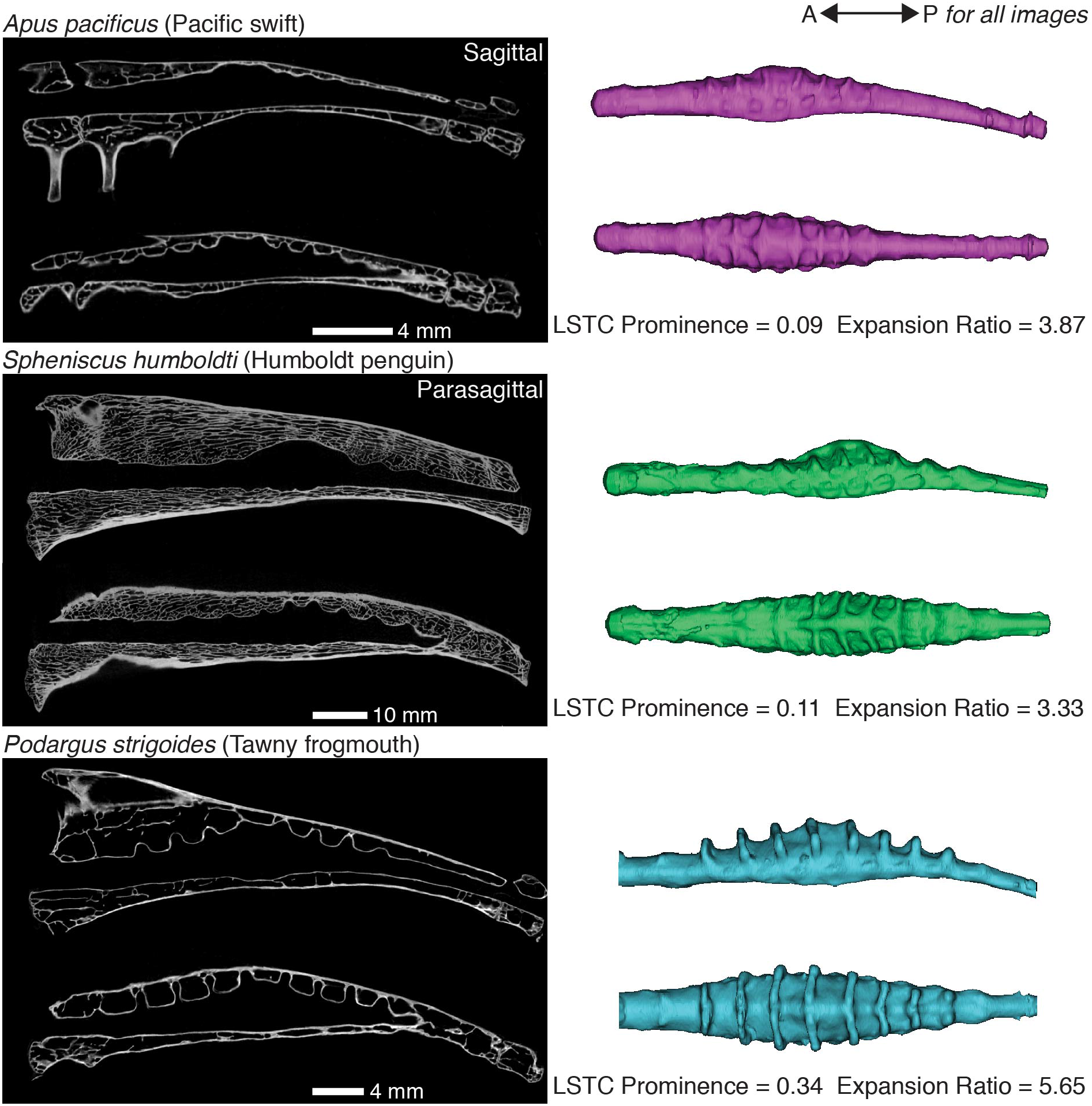
The morphology of the synsacral vertebral canal varies across bird species. These are sagittal and parasagittal sections from μ-CT scans of the synsacra of three bird species paired with lateral and dorsal views of the corresponding vertebral canal endocasts. Although *A. pacificus* and *S. humboldti* have similar values for the metrics we calculated, the shape of their endocasts differ, particularly in the orientation of the LSTCs (see dorsal views of endocasts). We calculated much higher values of both metrics for *P. strigoides* than for the other two species shown here, and the endocast of *P. strigoides* exhibits correspondingly visually prominent LSTCs and central expansion.

**Figure 5.**
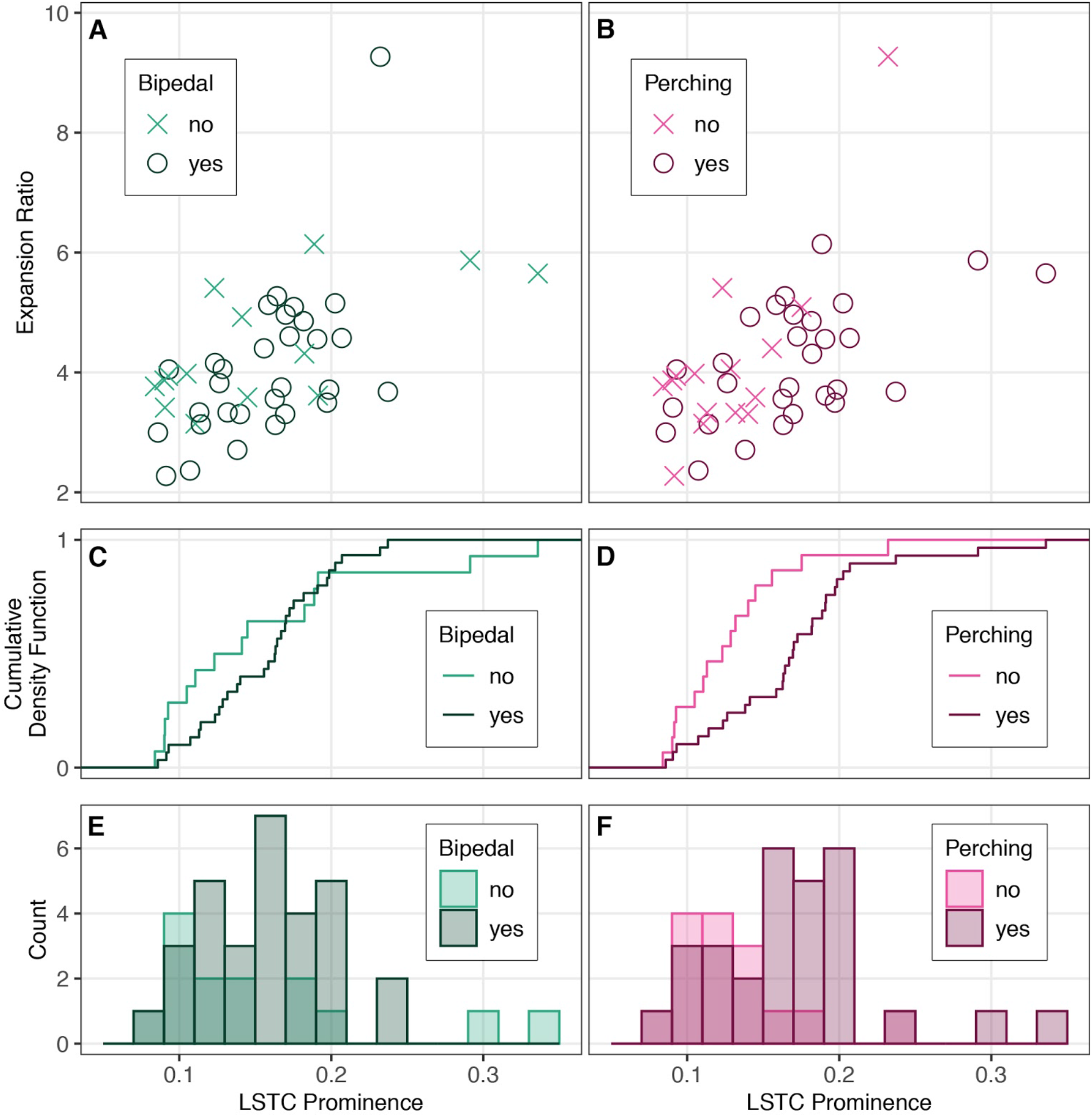
The LSTC prominence and expansion ratio of neural canal region of the LSO across 44 bird species does not clearly correspond with bipedalism, although there is a trend of increased canal prominence in birds that perch. The scatterplots in (A) and (B) form a morphospace based on canal prominence and expansion ratio. Each point represents one species, colored by whether or not they move bipedally or perch, respectively. (C) and (D) plot the corresponding cumulative density functions for LSTC prominence, while (E) and (F) are the corresponding histograms for LSTC prominence. Kolmogorov-Smirnov tests confirm a trend of more prominent LSTCs prominence in perching vs. non-perching species (D=0.556, p=0.0032) but not bipedal vs. non-bipedal (D=0.3, p=0.286).

The bird species in our sample occupy a large area of the morphospace defined by our computed metrics (Fig. 5; n=44; LSTC prominence: mean=0.156, median=0.157, st.dev.=0.054; expansion ratio: mean=4.173, median=3.904, st.dev.=1.204). In both of our locomotor mode comparisons, we found overlap between the two groups; in other words, the distributions of expansion ratios and LSTC prominences overlapped between birds that exhibit bipedal locomotion and those that are not bipedal, and between birds that perch and those that do not perch. Summary statistics for these locomotor groups are provided in Supplemental Table 1.

Thus, LSO osteological morphology does not clearly correspond with either perching or bipedal locomotor modes. However, we did find a trend toward more prominent LSTCs in birds that perch vs. birds that do not perch, as indicated by the shift in the cumulative density function and histogram plots (Fig. 5D, F). Kolmogorov-Smirnov tests confirmed this trend (LSTC prominence, perching vs. non-perching: D=0.556, p=0.0032; bipedal vs. non-bipedal: D=0.3, p=0.286). We did not find a similar trend for the expansion ratio (perching vs. non-perching: D=0.216, p=0.657; bipedal vs. non-bipedal: D=0.262, p=0.448). We also found that higher LSTC prominence values are convergent across the avian phylogeny (Fig. 6), suggesting that phylogeny alone does not explain the variation in LSO osteological morphology.

**Figure 6.**
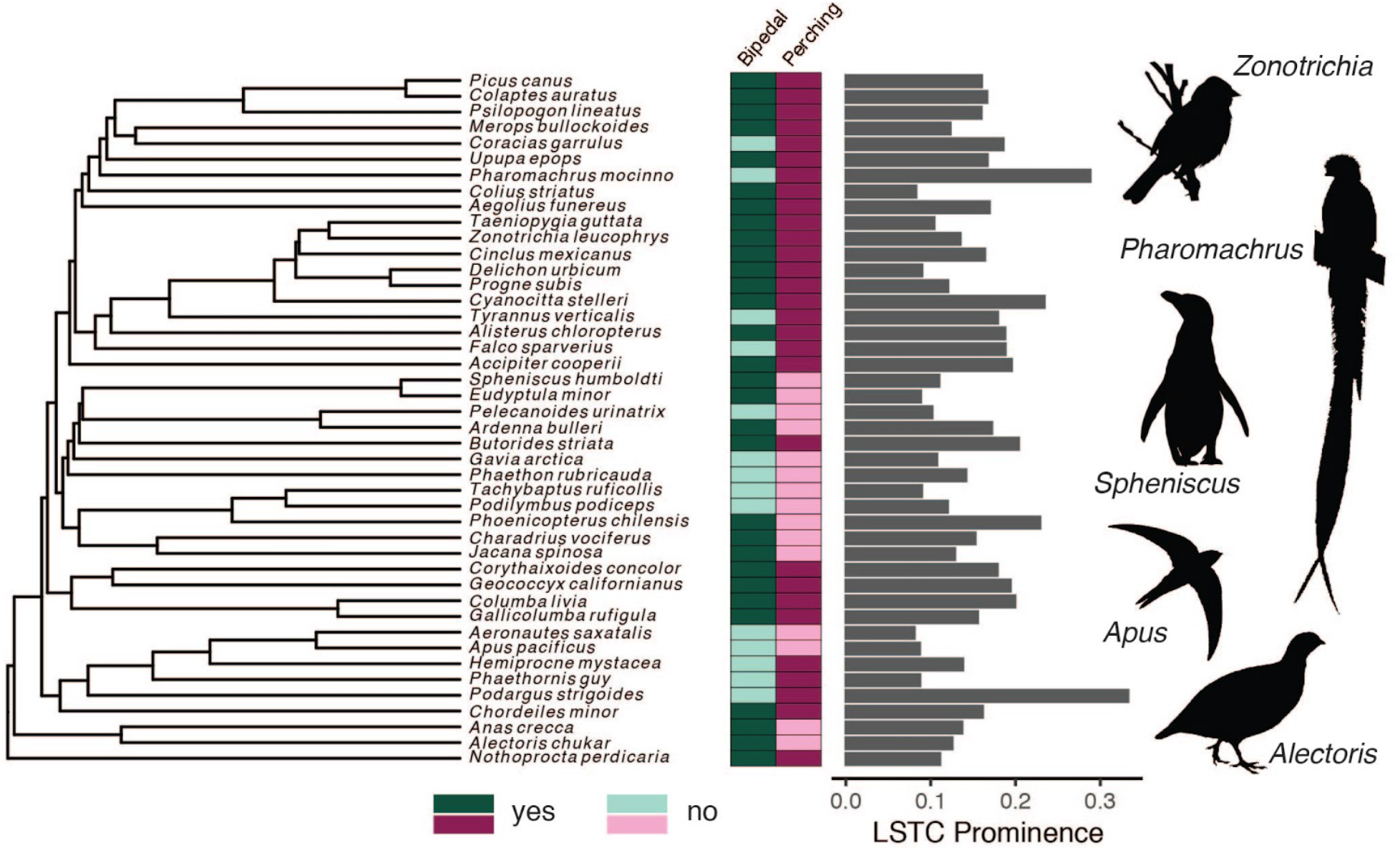
Locomotor mode and LSO osteological morphology are both convergent across Aves. Our interspecific sample is shown on a genus-level phylogeny from Cooney et al. (2017) based on Prum et al. (2015) and Jetz et al. (2012). Darker colors in the boxes to the right of the phylogeny indicate that the species perches or exhibits bipedal locomotion. Lighter colors indicate that the species does not perch or does not move bipedally. The computed LSTC prominence is shown on the bar chart.

For some bird species, we measured notably high values of our computed metrics. *Pharomachrus mocinno* (resplendent quetzal) and *P. strigoides* (tawny frogmouth) both exhibit more prominent LSTCs (Fig. 5 and Fig. 6) relative to the expansion in the neural canal. Both are perching species that rarely move about the ground bipedally; however, other species in our sample fit these criteria yet do not have such elaborated synsacral vertebral canal morphologies. *Phoenicopterus chilensis* (Chilean flamingo) exhibits an anomalously high expansion ratio. Flamingos are noteworthy not only for long legs and balancing on one foot, but also for their feeding habit, which requires them to lower and swing their head through the water while keeping their torso stable. Spoonbills exhibit a similar behavior (Rico-Guevara et al., 2019), and it would be interesting to compare their LSO morphology with that of flamingos. Perching and balancing on one foot may be the types of balance-intensive behaviors partially enabled by exaggerated LSO morphology.

Measurements of LSTC prominence and expansion ratio in species for which we had multiple samples demonstrate that intraspecific variation in LSO osteological morphology is reasonably small compared to interspecific variation, validating the use of these metrics in interspecific studies (Fig. 7). Intraspecific morphological consistency also supports a functional role for this morphology. One contrasting pair within these five species might be particularly illustrative of a locomotor ecology signal in LSO osteology. *A. pacificus*, the Pacific swift, exhibits the quintessentially aerial lifestyle of swifts (Apodidae) and is unable to perch or move bipedally. In contrast, *H. mystacea*, the moustached treeswift of the sister family Hemiprocnidae, has a modified tarsus such that it can perch. *A. pacificus* and *H. mystacea* occupy non-overlapping parts of this morphospace, with *H. mystacea* exhibiting more exaggerated morphological characteristics.

**Figure 7.**
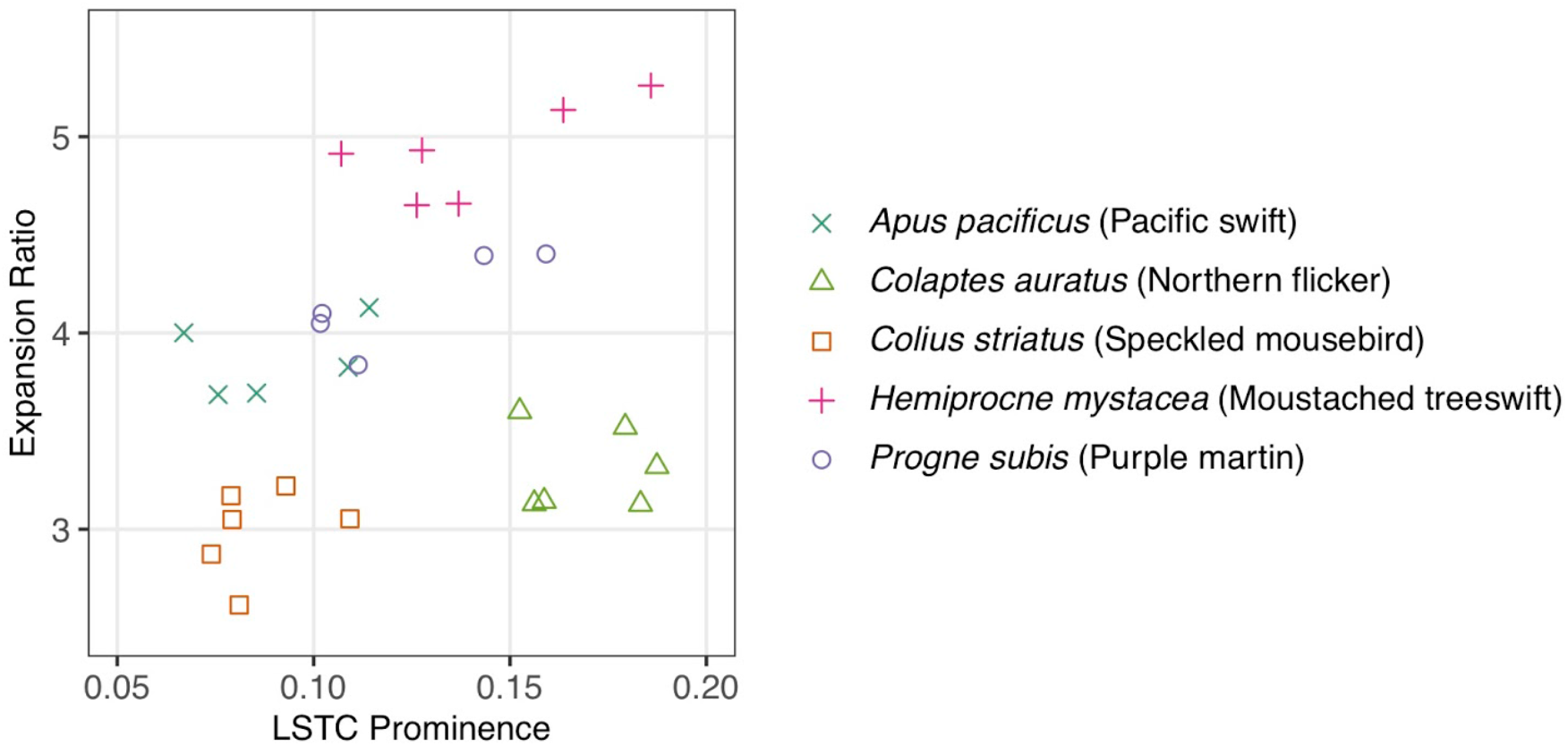
Multiple samples of five bird species occupy different areas of LSO osteological morphospace, indicating that interspecific variation in LSO osteological morphology is greater than intraspecific variation.

Overall, the metrics we developed for exploring LSO osteological morphology are useful for differentiating LSO morphotypes. We also found that the LSTC prominence metric was useful for understanding a possible relationship between locomotor ecology and the morphology of the LSO neural canal. We note again, however, the limitations of our sample size given the extreme diversity and number of avian species, the difficulty in assigning locomotor modes, and the lack of a phylogenetic analysis. In addition, our metrics are limited in their ability to fully describe the three-dimensional morphology of the vertebral canal space, such as the orientation and shape of the LSTCs. For instance, Jelgersma (1951) qualitatively described variation among species in the size of the bony gaps between LSTCs, which is not captured by either of our metrics. A future comparative analysis using 3D morphometric methods (e.g., alpha shapes, Gardiner et al., 2018; SPHARM, Styner et al., 2006 and Shen et al., 2009) may be an instructive ecomorphological investigation.

The soft tissue assessments using diceCT revealed the soft tissue morphology of the passerine LSO as similar to the morphology described for other bird species by previous researchers using histology and dissections (e.g., Necker, 2006). The method verified the presence of space for cerebrospinal fluid within the LSTCs and around the glycogen body (Fig. 2). Although immersion in I_2_KI solution can cause shrinkage, sections parallel to the sagittal and coronal planes in *Zonotrichia leucophrys* demonstrated that the spinal cord to the anterior and posterior of the LSO area almost completely fills the vertebral canal even after 11 days in solution, indicating minimal shrinkage in our preparation (Fig. 2c). In the scans of *Taeniopygia guttata*, the LSTCs and area around the glycogen body were free of soft tissue as in the *Zonotrichia leucophrys* scans, although there was potentially some shrinkage in the anterior portion of the spinal cord (Supplemental Fig. 1).

## Summary and Future Research

We constructed a comparative dataset that demonstrates the ubiquity of a specialized lumbosacral vertebral canal morphology across Aves, indicating that most—if not all—extant bird species possess an LSO. Based on a series of anatomical, physiological, and behavioral studies, Necker (2006) proposed that the LSO allows birds to sense body position distinctly from head position and thus perform two distinct locomotor functions: flying and “ground contact” (Rosenberg and Necker, 2002). Our quantitative comparative data allow us to refine the locomotor function aspect of this hypothesis. Specifically, we did not find a connection between a bipedal lifestyle and LSO osteological traits. We did find a trend toward more exaggerated LSO osteological traits in birds with a perching lifestyle, although the set of species that perch does overlap in trait space with the non-perching set.

We find Necker’s hypothesis that the LSO supports multiple, distinct locomotor modules compelling, and, our observations suggest the LSO may have a specific role in highly balance-intensive behaviors as evidenced by its exaggerated morphology in perching birds. Perching may be a locomotor behavior worthy of more detailed study in the context of the LSO, as might other hindlimb-based, balance-intensive, locomotor modes (e.g., balancing on one foot like flamingos). We note, though, that our classification of bipedalism encompasses a number of different locomotor modes, including hopping and alternating strides, and may not fully capture the bipedal performance measure most relevant to LSO function.

The existence of the LSO across Aves highlights several gaps in our understanding of avian neural and locomotor systems. Advancing understanding of LSO function will require simultaneous investigations at the organismal, organ, and cellular levels. One limitation of this current study is our categorization of locomotor lifestyle. Because most birds are capable of performing many locomotor functions, a binary categorization of most prominent locomotor traits of a species based on descriptive data is ultimately subjective. The development of robust, quantitative metrics for determining balance performance—ideally measured in a variety of taxa across Aves—would greatly assist future studies of avian locomotor function, including phylogenetic hypothesis tests of the impact of the LSO on organismal function. Gaining a detailed understanding of how the avian body moves relative to the head in a variety of locomotor maneuvers may also be insightful, as the LSO may not specifically enable ground-based locomotion. In flight and on the ground, birds maintain exquisite head stability while the body accelerates (Erichsen et al. 1989, Katzir et al., 2001, Warrick et al., 2002).

In addition to its putative role in locomotor behavior, perhaps the most conspicuous gap in our understanding of the LSO is the lack of experimental evidence for proximal mechanisms, in particular how it may transduce and encode acceleration. Fortunately, experimental techniques have advanced considerably since the bulk of published LSO research was completed. After discovering the LSTCs, Necker submitted that the LSTCs and the accessory lobes are the lumbosacral equivalent of the semicircular canals of the inner ear, acting to sense rotational motion while the dentate ligament stabilizes the spinal cord (as described in Necker, 2006, and citations therein). However, this explanation fails to account for the presence of the glycogen body, and the soft tissue elements of the LSO may act in concert to sense acceleration forces (as has also been noted in abstracts by Wold & Daley, 2019 and Kamska et al., 2020).

We suggest that the accessory lobes, LSTCs, and cerebrospinal fluid might act as a primarily rotational accelerometer while the motion of the glycogen body relative to that of the spinal cord acts as a translational accelerometer. Although mechanosensory neurons have been identified in the vertebrate spinal cord and shown to have locomotor function (Agduhr, 1922; Wyart et al., 2009), the avian spinal cord at the LSO remains mysterious. Determining the locations and identities of mechanosensory cells of the spinal cord and their responses to the relevant mechanical stimuli, including cerebrospinal fluid motion and tissue vibration, will help distinguish these hypotheses. Further, the putative neural circuitry of the LSO may function locally within the spinal cord and/or project to the cerebellum, which is known to receive balance information from the inner ear and contribute to calibration and coordination of movement (Kandel et al., 2000). It is important, however, to note that all hypotheses of LSO function lack conclusive evidence, and many should still be considered until some of these knowledge gaps are filled.

It would be difficult to understate the implications of elucidating LSO function for understanding the evolution of extant birds. Most birds spend a minority of their time in the air, thus legs and their use are an essential component of modern avian morphological and ecological diversity (Zeffer et al., 2002; Abourachid and Höfling, 2012). Gatesy and Dial (1996) proposed that the evolution of multiple “locomotor modules” (i.e., an aerial, forelimb-based module that is decoupled from the ancestral hindlimb-based module) contributed to the ecological diversity and evolutionary success of modern Aves, in part by enabling the diversification of avian hindlimbs (Gatesy and Dial, 1996; Gatesy and Middleton, 1997). The hypothesized role of the LSO as a secondary balance organ would increase neural modularity and allow distinct but coordinated stabilization of the avian body and the head, facilitating the evolution of the multiple locomotor modules proposed by Gatesy and Dial (1996). A curious analogy can be made with gyroscopic sensing in flying insects, which also exhibit superb head stabilization behavior. Flying insects have multiple organs that support gyroscopic sensing and balance: the wings and/or halteres at the thorax, and the antennae on the head (Rauscher and Fox, 2018; Sane et al., 2007; Dickerson et al., 2014 and 2019). If a mechanosensory function for the LSO is conclusively demonstrated, then it could be a key innovation for Aves as well as a trait that enabled the evolution of flight and/or exquisite bipedal performance, including perching.

## Supporting information

All supplemental files (data and code)

## Data Accessibility

Raw LSTC area data, Matlab code, calculated metric data for creating figures, and R code are provided as supplemental files.

## Acknowledgements

The authors thank the Ornithology Collection at the Burke Museum of Natural History and Culture and the Division of Birds at the Smithsonian National Museum of Natural History for access to specimens. Sharlene Santana, Adam Summers, and Michelle Hickner provided access and help with CT scanning. Cassandra Fieldson and Calvin Davis produced endocasts. Kimberly Miller helped with soft tissue anatomical investigations, and Tanvi Deora provided helpful discussion and references on insect inertial sensing. Alejandro Rico-Guevara and Tom Daniel provided helpful comments and discussion on a draft of the manuscript.

## Supplemental Information

**Supplemental Table 1.** Summary statistics for the LSTC prominence and expansion ratio for the entire sample and for subsamples defined by locomotor ecology.

|  | Mean | Median | Standard deviation |
| --- | --- | --- | --- |
| **All (n=44)** |  |  |  |
| LSTC prominence | 0.156 | 0.157 | 0.054 |
| Expansion ratio | 4.173 | 3.904 | 1.204 |
| **Perching (n=29)** |  |  |  |
| LSTC prominence | 0.171 | 0.170 | 0.055 |
| Expansion ratio | 4.168 | 4.05 | 0.971 |
| **Not perching (n=15)** |  |  |  |
| LSTC prominence | 0.128 | 0.123 | 0.039 |
| Expansion ratio | 4.183 | 3.867 | 1.602 |
| **Bipedal (n=30)** |  |  |  |
| LSTC prominence | 0.157 | 0.163 | 0.04 |
| Expansion ratio | 4.067 | 3.786 | 1.291 |
| **Not bipedal (n=14)** |  |  |  |
| LSTC prominence | 0.155 | 0.132 | 0.077 |
| Expansion ratio | 4.401 | 3.96 | 0.998 |

**Supplemental Figure 1.**
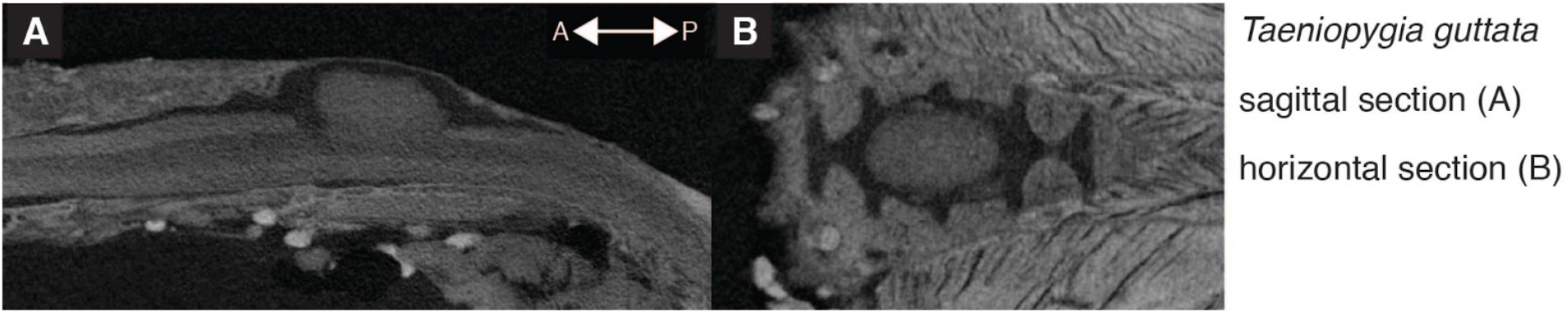
Contrast-enhanced CT scan of the lumbosacral organ (LSO) of Taeniopygia guttata demonstrates fluid space with the lumbosacral transverse canals (LSTCs).

## Supplemental Information - List of Museum Specimens

Acronyms: UWBM: Ornithology Collection, Burke Museum of Natural History and Culture; USNM: Division of Birds, Smithsonian National Museum of Natural History

*Accipiter cooperii* UWBM 65054

*Aegolius funereus* UWBM 44737

*Aeronautes saxatalis* UWBM 34156

*Alectoris chukar* UWBM 19808

*Alisterus chloropterus* UWBM 43544

*Anas crecca* UWBM 59245

*Apus pacificus* UWBM 44398, UWBM 44399, UWBM 46954, UWBM 46955, UWBM 46956

*Butorides striatus* UWBM 45810

*Charadrius vociferus* UWBM 88808

*Cinclus mexicanus* UWBM 50254

*Colaptes auratus* UWBM 19131, UWBM 32004, UWBM 32500, UWBM 87743, USNM 553938, USNM 610529

*Colius striatus* UWBM 38252, UWBM 38253, UWBM 52742, USNM 428106, USNM 491986,

USNM 558540

*Columba livia* UWBM 76160

*Coracia garrulus* UWBM 56647

*Cordeiles minor* UWBM 65014

*Corythaixoides concolor* UWBM 52910

*Cyanocitta stelleri* UWBM 52505

*Delichon urbica* UWBM 47240

*Eudyptula minor* UWBM 57504

*Falco sparverius* UWBM 26276

*Gallicolumba rufigula* UWBM 43040

*Gavia arctica* UWBM 28567

*Geococcyx californicus* UWBM 34139

*Hemiprocne mystacea* UWBM 58752, UWBM 60348, UWBM 62344, USNM 559041, USNM

560829, USNM 572397

*Jacana spinosa* UWBM 21277

*Megalaima lineata* UWBM 64902

*Merops bullockoides* UWBM 57056

*Nothoprocta perdicaria* UWBM 65156

*Pelecanoides urinator* UWBM 60504

*Phaethon rubricauda* UWBM 68941

*Pharomachus moccino* UWBM 18395

*Phaethornis guy* UWBM 22394

*Phoenicopterus chilensis* UWBM 39909

*Picus canus* UWBM 51782

*Podargus strigoides* UWBM 76160

*Podilymbus podiceps* UWBM 30133

*Progne subis* UWBM 45892, UWBM 55860, USNM 501361, USNM 556284, USNM 610520

*Puffinus bulleri* UWBM 55371

*Spheniscus humboldti* UWBM 91277

*Tachybaptus ruficollis* UWBM 86353

*Taeniopygia guttata* (from laboratory colony)

*Tyrannus verticalis* UWBM 35770

*Upupa epops* UWBM 46408

*Zonotrichia leucophrys* (from laboratory colony)

