## Supplementary material for "The balance hypothesis for the avian lumbosacral organ and an exploration of its morphological variation": All supplemental files (data and code): UW_LSOteam_Apr202biorxiv_LSOmetrics.m

```
% Stanchak, French, Perkel, Brunton 2020
%
% Calculate SOR (standard deviation of residuals) and ExRat (expansion
% ratio of LSO) of cross-sectional areas of synsacral CT scans
% See manuscript for details of data collection.

% Read in data file
data = readtable('LSO_Area_Data_012320.csv');

% Set up output file
ndata = width(data)-1;
LSOdata = table('Size', [ndata 3], 'VariableTypes', {'double', 'double', 'double'});
LSOdata.Properties.VariableNames = {'SOR', 'SORall', 'ExRat'};
LSOdata.Properties.RowNames = data.Properties.VariableNames(2:end);

% Calculate ExRat & SOR
n = width(data);

for i =2:n
    sp = data{1:end, i};
    
    % get rid of nan
    sp(~any(~isnan(sp), 2),:)=[];
    
    last = sp(end);
    first = sp(1);
    a = [first last];
    [middle,I] = max(sp);
    
    % SOR calculated around max
    SORdata = sp(I-5:I+5);
    runMean = movmean(SORdata, 3);
    resids = SORdata./runMean;
    SOR = std(resids);
    
    % SOR calculated with all areas
    runMeanall = movmean(sp, 3);
    residsall = sp./runMeanall;
    SORall = std(residsall);
    
    ExRat = max(runMean)/mean(a);
    
    LSOdata(i-1,3) = {ExRat};
    LSOdata(i-1,1) = {SOR};
    LSOdata(i-1,2) = {SORall};
end

writetable(LSOdata, 'LSOdata.csv', 'WriteRowNames', true)

    
% code for generating example data for Fig 3    

data1 = data{1:end, 55}
data1(~any(~isnan(data1), 2),:)=[]
last=data1(end)
first=data1(1)
a=[first last]
[middle, I]=max(data1)
SORdata = data1(I-5:I+5)
runMean=movmean(SORdata, 3)
resids = SORdata./runMean
D = [SORdata, runMean, resids]
writematrix(D, 'plotdata.csv')
```
