## Supplementary material for "The balance hypothesis for the avian lumbosacral organ and an exploration of its morphological variation": All supplemental files (data and code): UW_LSOteam_Apr2020biorxiv_FigsStats.R

```
# Stanchak, French, Perkel, Brunton 2020

#set wd

library(ape)
library(geiger)
library(ggplot2)
library(ggtree)
library(treeio)
library(dplyr)
library(cowplot)
library(egg)
library(extrafont)
#loadfonts(device = "pdf", quiet = FALSE)

# read in tree with changed names (based on Prum_merge_Genera_2016_04_22_Hackett_Stage2_10K_MCC.newick.tre 
# from Cooney et al. 2017)
LSOtree<-read.nexus(file="LSO_taxa.nex")

data<-read.csv("LSOdata_021020.csv")

Hemiprocne <- data[which(data$Epithet == "Hemiprocne_mystacea"),]
Apus <- data[which(data$Epithet == "Apus_pacificus"),]
Colius <- data[which(data$Epithet == "Colius_striatus"),]
Colaptes <- data[which(data$Epithet == "Colaptes_auratus"),]
Progne <- data[which(data$Epithet == "Progne_subis"),]

Hemiprocne_mean_SOR <- mean(Hemiprocne$SOR)
Hemiprocne_mean_SORall <- mean(Hemiprocne$SORall)
Hemiprocne_mean_ExRat <- mean(Hemiprocne$ExRat)
Hemiprocne_mean <- as.data.frame(cbind("Hemiprocne", "mystacea", "Hemiprocne mystacea", "Hemiprocne_mystacea", "Hemiprocne_mystacea_mean", Hemiprocne_mean_SOR,
                                       Hemiprocne_mean_SORall, Hemiprocne_mean_ExRat, paste(Hemiprocne$Fly[1]), paste(Hemiprocne$Perching[1]),
                                       paste(Hemiprocne$Float[1]), paste(Hemiprocne$Dive[1]), paste(Hemiprocne$Bipedal[1]),
                                       paste(Hemiprocne$Perch_Tree[1]), paste(Hemiprocne$Bipedal_Tree[1])))
names(Hemiprocne_mean) <- names(data)

Apus_mean_SOR <- mean(Apus$SOR)
Apus_mean_SORall <- mean(Apus$SORall)
Apus_mean_ExRat <- mean(Apus$ExRat)
Apus_mean <- as.data.frame(cbind("Apus", "pacificus", "Apus pacificus", "Apus_pacificus", "Apus_pacificus_mean", Apus_mean_SOR,
                                 Apus_mean_SORall, Apus_mean_ExRat, paste(Apus$Fly[1]), paste(Apus$Perching[1]),
                                 paste(Apus$Float[1]), paste(Apus$Dive[1]), paste(Apus$Bipedal[1]),
                                 paste(Apus$Perch_Tree[1]), paste(Apus$Bipedal_Tree[1])))
names(Apus_mean) <- names(data)

Colius_mean_SOR <- mean(Colius$SOR)
Colius_mean_SORall <- mean(Colius$SORall)
Colius_mean_ExRat <- mean(Colius$ExRat)
Colius_mean <- as.data.frame(cbind("Colius", "striatus", "Colius striatus", "Colius_striatus", "Colius_striatus_mean", Colius_mean_SOR,
                                   Colius_mean_SORall, Colius_mean_ExRat, paste(Colius$Fly[1]), paste(Colius$Perching[1]),
                                   paste(Colius$Float[1]), paste(Colius$Dive[1]), paste(Colius$Bipedal[1]),
                                   paste(Colius$Perch_Tree[1]), paste(Colius$Bipedal_Tree[1])))
names(Colius_mean) <- names(data)

Colaptes_mean_SOR <- mean(Colaptes$SOR)
Colaptes_mean_SORall <- mean(Colaptes$SORall)
Colaptes_mean_ExRat <- mean(Colaptes$ExRat)
Colaptes_mean <- as.data.frame(cbind("Colaptes", "auratus", "Colaptes auratus", "Colaptes_auratus", "Colaptes_auratus_mean", Colaptes_mean_SOR,
                                     Colaptes_mean_SORall, Colaptes_mean_ExRat, paste(Colaptes$Fly[1]), paste(Colaptes$Perching[1]),
                                     paste(Colaptes$Float[1]), paste(Colaptes$Dive[1]), paste(Colaptes$Bipedal[1]),
                                     paste(Colaptes$Perch_Tree[1]), paste(Colaptes$Bipedal_Tree[1])))
names(Colaptes_mean) <- names(data)

Progne_mean_SOR <- mean(Progne$SOR)
Progne_mean_SORall <- mean(Progne$SORall)
Progne_mean_ExRat <- mean(Progne$ExRat)
Progne_mean <- as.data.frame(cbind("Progne", "subis", "Progne subis", "Progne_subis", "Progne_subis_mean", Progne_mean_SOR,
                                   Progne_mean_SORall, Progne_mean_ExRat, paste(Progne$Fly[1]), paste(Progne$Perching[1]),
                                   paste(Progne$Float[1]), paste(Progne$Dive[1]), paste(Progne$Bipedal[1]),
                                   paste(Progne$Perch_Tree[1]), paste(Progne$Bipedal_Tree[1])))
names(Progne_mean) <- names(data)

newdata5 <- data[which(data$Epithet != "Hemiprocne_mystacea"),]
newdata4 <- newdata5[which(newdata5$Epithet != "Apus_pacificus"),]
newdata3 <- newdata4[which(newdata4$Epithet != "Colius_striatus"),]
newdata2 <- newdata3[which(newdata3$Epithet != "Colaptes_auratus"),]
newdata1 <- newdata2[which(newdata2$Epithet != "Progne_subis"),]

newdata <- rbind(newdata1, Hemiprocne_mean, Apus_mean, Colius_mean, Colaptes_mean, Progne_mean)
row.names(newdata)<-newdata$Epithet
fivedata = rbind(Hemiprocne, Apus, Colius, Colaptes, Progne)

# Kolmorgorov-Smirnov tests

perchdata <- newdata[which(newdata$Perching == "yes"),]
noperchdata <- newdata[which(newdata$Perching == "no"),]
ks.test(as.numeric(perchdata$SOR), as.numeric(noperchdata$SOR))
ks.test(as.numeric(perchdata$ExRat), as.numeric(noperchdata$ExRat))

bipedaldata <- newdata[which(newdata$Bipedal == "yes"),]
nobipedaldata <- newdata[which(newdata$Bipedal == "no"),]
ks.test(as.numeric(bipedaldata$SOR), as.numeric(nobipedaldata$SOR))
ks.test(as.numeric(bipedaldata$ExRat), as.numeric(nobipedaldata$ExRat))

# Summary Stats

mean_SOR_all <- mean(as.numeric(newdata$SOR))
median_SOR_all <- median(as.numeric(newdata$SOR))
sd_SOR_all <- sd(as.numeric(newdata$SOR))
mean_exrat_all <- mean(as.numeric(newdata$ExRat))
median_exrat_all <- median(as.numeric(newdata$ExRat))
sd_exrat_all <- sd(as.numeric(newdata$ExRat))

mean_SOR_perch <- mean(as.numeric(perchdata$SOR))
median_SOR_perch <- median(as.numeric(perchdata$SOR))
sd_SOR_perch <- sd(as.numeric(perchdata$SOR))
mean_exrat_perch <- mean(as.numeric(perchdata$ExRat))
median_exrat_perch <- median(as.numeric(perchdata$ExRat))
sd_exrat_perch <- sd(as.numeric(perchdata$ExRat))

mean_SOR_noperch <- mean(as.numeric(noperchdata$SOR))
median_SOR_noperch <- median(as.numeric(noperchdata$SOR))
sd_SOR_noperch <- sd(as.numeric(noperchdata$SOR))
mean_exrat_noperch <- mean(as.numeric(noperchdata$ExRat))
median_exrat_noperch <- median(as.numeric(noperchdata$ExRat))
sd_exrat_noperch <- sd(as.numeric(noperchdata$ExRat))

mean_SOR_bipedal <- mean(as.numeric(bipedaldata$SOR))
median_SOR_bipedal <- median(as.numeric(bipedaldata$SOR))
sd_SOR_bipedal <- sd(as.numeric(bipedaldata$SOR))
mean_exrat_bipedal <- mean(as.numeric(bipedaldata$ExRat))
median_exrat_bipedal <- median(as.numeric(bipedaldata$ExRat))
sd_exrat_bipedal <- sd(as.numeric(bipedaldata$ExRat))

mean_SOR_nobipedal <- mean(as.numeric(nobipedaldata$SOR))
median_SOR_nobipedal <- median(as.numeric(nobipedaldata$SOR))
sd_SOR_nobipedal <- sd(as.numeric(nobipedaldata$SOR))
mean_exrat_nobipedal <- mean(as.numeric(nobipedaldata$ExRat))
median_exrat_nobipedal <- median(as.numeric(nobipedaldata$ExRat))
sd_exrat_nobipedal <- sd(as.numeric(nobipedaldata$ExRat))

# Tree plots

trimmed<-geiger::treedata(LSOtree, newdata, sort=T)
trim.data<-trimmed$data
trim.tree<-trimmed$phy
name.check(trim.tree, trim.data)
plot(trim.tree)

LSOtree<-rename_taxa(trim.tree, as.data.frame(trim.data), Epithet, Binomial) 
Locomotion <- cbind(paste(newdata$Bipedal_Tree), 
                    paste(newdata$Perch_Tree)) #, paste(newdata$Float), paste(newdata$Dive))
rownames(Locomotion)<-newdata$Binomial
colnames(Locomotion)<-as.character(c("Bipedal", "Perching")) #, "Swim", "Dive"))
Locomotion<-as.data.frame(Locomotion)

trim.data.df<-as.data.frame(trim.data)
df<-data.frame(label = LSOtree$tip.label, genus = trim.data.df$Genus, 
               species = trim.data.df$specific)
temp <- ggtree(LSOtree) %<+% df + xlim(0,140) + ylim(0,47) + 
  geom_tiplab(aes(label=paste0('italic(', genus,')~italic(', species,')')), 
              parse=T, size=2, family="Helvetica", offset=1) +
  theme(plot.margin = unit(c(0, 0, 0, 0), "cm"))
temp

heat<-gheatmap(temp, Locomotion, offset=40, width=0.2, font.size=2.5, colnames_position = "top",
               colnames_angle=50, hjust=0, color="black") +
  scale_fill_manual(breaks=c("noP", "yesP", "noB", "yesB"), values=c("#a4d8c9", "#f5a9d0", "#0e4f3c", "#8b1953")) + 
  theme(legend.position="none") #+
heat

d <- filter(temp, isTip) %>% select(c(label, y))
colnames(d)<-c("Binomial", "y")

dd1 <- left_join(newdata, d, by="Binomial")

p1 <- ggplot(dd1, aes(y, as.numeric(SOR))) + geom_col() + 
  coord_flip() + theme_tree2() + theme(legend.position='none') +
  theme(text=element_text(size=10, family="Helvetica")) +
  ylab("LSTC Prominence") + ylim2(temp) +
  theme(plot.margin = unit(c(0, 0, 0, 0), "cm"))
p1 

treegrid <- plot_grid(heat, p1, ncol=2, rel_widths = c(3, 1), align="h",
                      labels=NULL)
treegrid

ggsave("treetemp.pdf",plot=treegrid, width = 160, height = 110, dpi=300, units="mm") 

# Scatterplot of just 5 taxa

p<-ggplot(data=fivedata, aes(x=as.numeric(SOR), y=as.numeric(ExRat))) + 
  geom_point(aes(color=Binomial, shape=Binomial), size=2) + 
  scale_color_manual(values=c("#1b9e77", "#66a61e", "#d95f02", "#e7298a", "#7570b3"),
                     labels=c(bquote(paste(italic("Apus pacificus "), "(Pacific swift)")), 
                              bquote(paste(italic("Colaptes auratus "), "(Northern flicker)")),
                              bquote(paste(italic("Colius striatus "), "(Speckled mousebird)")),
                              bquote(paste(italic("Hemiprocne mystacea "), "(Moustached treeswift)")),
                              bquote(paste(italic("Progne subis "), "(Purple martin)")))) + 
  scale_shape_manual(values=c(4, 2, 0, 3, 1),
                     labels=c(bquote(paste(italic("Apus pacificus "), "(Pacific swift)")), 
                              bquote(paste(italic("Colaptes auratus "), "(Northern flicker)")),
                              bquote(paste(italic("Colius striatus "), "(Speckled mousebird)")),
                              bquote(paste(italic("Hemiprocne mystacea "), "(Moustached treeswift)")),
                              bquote(paste(italic("Progne subis "), "(Purple martin)")))) +
  theme_bw() + theme(panel.grid.minor=element_blank()) +
  theme(aspect.ratio=1, text=element_text(size=10, family="Helvetica")) +
  theme(legend.text=element_text(size = 10, family = "Helvetica", face="italic")) +
  theme(axis.text=element_text(size=10)) +
  xlab("LSTC Prominence") + ylab("Expansion Ratio") + theme(legend.title=element_blank()) +
  xlim(0.05,0.2) + ylim(2.5,5.5)
p
ggsave("Fig6.pdf",plot=p, width = 180, height = 80, dpi=300, units="mm") 

# Scatterplots and histograms of all taxa for bipedal/perching

g_bipedal<-ggplot(newdata, aes(x = as.numeric(SOR), y = as.numeric(ExRat))) + 
  geom_point(data=newdata, aes(color=Bipedal, shape=Bipedal), size=3.5) + 
  theme_bw() + theme(panel.grid.minor=element_blank()) +
  scale_color_manual(values=c("#32a885", "#0b3f30")) + 
  theme(aspect.ratio=1) + theme(text=element_text(size=10, family="Helvetica")) +
  theme(legend.position=c(0.2,0.75), legend.text=element_text(size=10, family="Helvetica"),
        legend.box.background = element_rect(colour = "black")) +
  scale_shape_manual(values = c(4, 1)) + 
  xlim(0.05,0.35) + ylim(2.2,9.8) +
  theme(axis.title.x=element_blank(), axis.text.x=element_blank(), 
        axis.ticks.x=element_blank(), axis.text.y=element_text(size=10)) +
  ylab("Expansion Ratio") + theme(plot.margin = unit(c(3, 0, 0, 5.5),"mm"))
g_bipedal

g_perch<-ggplot(newdata, aes(x = as.numeric(SOR), y = as.numeric(ExRat))) + 
  geom_point(data=newdata, aes(color=Perching, shape=Perching), size=3.5) + 
  theme_bw() + theme(panel.grid.minor=element_blank()) +
  scale_color_manual(values=c("#ec54a1", "#741545")) + 
  theme(aspect.ratio=1) + theme(text=element_text(size=10, family="Helvetica")) +
  theme(legend.position=c(0.2,0.75), legend.text=element_text(size=10, family="Helvetica"),
        legend.box.background = element_rect(colour = "black")) +
  scale_shape_manual(values = c(4,1)) + xlim(0.05,0.35) + ylim(2.2,9.8) +
  theme(axis.title.x=element_blank(), axis.text.x=element_blank(), axis.ticks.x=element_blank(),
        axis.title.y=element_blank(), axis.text.y=element_blank(), axis.ticks.y=element_blank()) +
  ylab("Expansion Ratio") + theme(plot.margin = unit(c(3, 5.5, 0, 0),"mm"))
g_perch

h_bipedalSOR <- ggplot(newdata, aes(x=as.numeric(SOR), fill=Bipedal, color=Bipedal)) + 
  theme_bw() + theme(panel.grid.minor=element_blank()) + theme(aspect.ratio=0.5) +
  theme(text=element_text(size=10, family="Helvetica")) +
  geom_histogram(binwidth=0.02, alpha=0.3, position="identity") + 
  scale_fill_manual(values=c("#32a885", "#0b3f30")) +
  scale_color_manual(values=c("#32a885", "#0b3f30")) +
  xlim(0.05, 0.35) + ylim(0, 7) +
  xlab("LSTC Prominence") + ylab("Count") +
  theme(axis.text=element_text(size=10))+
  #theme(legend.position="none") +
  theme(legend.position=c(0.8,0.65), legend.text=element_text(size=10, family="Helvetica"),
        legend.box.background = element_rect(colour = "black")) +
  theme(plot.margin = unit(c(3, 0, 2, 5.5),"mm"))
h_bipedalSOR

h_perchSOR <- ggplot(newdata, aes(x=as.numeric(SOR), fill=Perching, color=Perching)) + 
  theme_bw() + theme(panel.grid.minor=element_blank()) + theme(aspect.ratio=0.5) +
  theme(text=element_text(size=10, family="Helvetica")) +
  geom_histogram(binwidth=0.02, alpha=0.3, position="identity") + 
  scale_fill_manual(values=c("#ec54a1", "#741545")) +
  scale_color_manual(values=c("#ec54a1", "#741545")) +
  xlim(0.05, 0.35) + ylim(0, 7) +
  theme(axis.title.y=element_blank(), axis.text.y=element_blank(), axis.ticks.y=element_blank(),
        axis.text.x=element_text(size=10)) +
  xlab("LSTC Prominence") + ylab("Count") +
  #theme(legend.position="none") +
  theme(legend.position=c(0.8, 0.65), legend.text=element_text(size=10, family="Helvetica"),
        legend.box.background = element_rect(colour = "black"))+
  theme(plot.margin = unit(c(3, 5.5, 2, 0),"mm"))
h_perchSOR

c_perchSOR <- ggplot(newdata, aes(x=as.numeric(SOR), color=Perching)) +stat_ecdf(show.legend=T) +
  theme_bw() + theme(panel.grid.minor=element_blank()) +
  theme(text=element_text(size=10, family="Helvetica")) +
  theme(legend.position="right") + ylab(NULL) + 
  scale_color_manual(values=c("#ec54a1", "#741545")) +
  xlim(0.05, 0.35) +
  theme(axis.title.x=element_blank(), axis.text.x=element_blank(), axis.ticks.x=element_blank(),
        axis.title.y=element_blank(), axis.text.y=element_blank(), axis.ticks.y=element_blank()) +
  scale_y_continuous(limits=c(0,1), breaks=c(0,1)) +
  theme(aspect.ratio=0.5) + 
  #theme(legend.position="none") +
  theme(legend.position=c(0.8, 0.4), legend.text=element_text(size=10, family="Helvetica"),
        legend.box.background = element_rect(colour = "black")) +
  theme(plot.margin = unit(c(3, 5.5, 0, 0), "mm"))
c_perchSOR

c_bipedalSOR <- ggplot(newdata, aes(x=as.numeric(SOR), color=Bipedal)) + 
  stat_ecdf(show.legend=T) +
  theme_bw() + theme(panel.grid.minor=element_blank()) +
  theme(text=element_text(size=10, family="Helvetica")) +
  theme(legend.position="right") +  ylab(NULL) + 
  scale_color_manual(values=c("#32a885", "#0b3f30")) +
  theme(axis.title.x=element_blank(), axis.text.x=element_blank(), axis.ticks.x=element_blank(),
        axis.text.y=element_text(size=10)) +
  theme(aspect.ratio=0.5) + ylab("Cumulative\nDensity Function") +
  scale_y_continuous(limits=c(0,1), breaks=c(0,1)) +
  xlim(0.05, 0.35) +
  #theme(legend.position="none") +
  theme(legend.position=c(0.8, 0.4), legend.text=element_text(size=10, family="Helvetica"),
        legend.box.background = element_rect(colour = "black")) + 
  theme(plot.margin = unit(c(3, 0, 0, 5.5), "mm"))
c_bipedalSOR

arranged<-ggarrange(g_bipedal, g_perch, c_bipedalSOR, c_perchSOR, h_bipedalSOR, h_perchSOR, ncol=2)

ggsave("Fig4.pdf",plot=arranged, width = 180, height = 220, dpi=300, units="mm") 

# Sample plot of areas and ratios

plotdata<-read.csv("plotdata_forR.csv", header=FALSE)

g_areas1<-ggplot(plotdata, aes(x=as.numeric(V4), as.numeric(V1))) + theme_classic() +
  theme(text=element_text(size=10, family="Helvetica")) + theme(aspect.ratio=0.6) +
  theme(axis.line=element_line(size=0.5)) +
  geom_line(size=0.5)+
  geom_line(aes(x=as.numeric(V4), as.numeric(V2)), linetype="dashed", size=0.5) +
  geom_point(aes(color=V5), shape=16, size=2) +
  scale_color_manual(values=c("#53863d", "#94c162")) +
  theme(legend.position="none")  + 
  scale_x_continuous(breaks=c(1, 3, 5, 7, 9, 11)) +
  # comment out y label below to produce properly sized graph then reinsert in photoshop
  labs(#y=expression(atop(paste("Cross sectional Area, A"[n]), "(Pixels)")), 
    x=expression("Cross section Index, n"))
g_areas1

ggsave("areas.pdf",plot=g_areas1, width = 80, height = 60, dpi=300, units="mm") 

g_ratios<-ggplot(plotdata, aes(x=as.numeric(V4), as.numeric(V3))) + theme_classic() +
  theme(text=element_text(size=10, family="Helvetica")) + theme(aspect.ratio=0.6) +
  theme(axis.line=element_line(size=0.5)) +
  geom_line(size=0.5)+
  geom_point(aes(color=V5), shape=16, size=2) +
  scale_color_manual(values=c("#53863d", "#94c162")) +
  geom_segment(aes(x=1, y=1, xend=11, yend=1), linetype="dashed", size=0.5) +
  #geom_hline(yintercept=1, linetype="dashed") +
  theme(legend.position="none") + scale_y_continuous(breaks=c(0.5, 0.75, 1, 1.25, 1.5)) + 
  scale_x_continuous(breaks=c(1, 3, 5, 7, 9, 11)) +
  labs(y=expression(paste("Normalized Area Ratio, R"[n])), 
       x=expression(paste("Cross section Index, n")))
g_ratios

ggsave("ratios.pdf",plot=g_ratios, width = 80, height = 60, dpi=300, units="mm") 
```
